# Present and future suitability of invasive and urban vectors through an environmentally-driven Mosquito Reproduction Number

**DOI:** 10.1101/2024.05.31.596775

**Authors:** Marta Pardo-Araujo, Roger Eritja, David Alonso, Frederic Bartumeus

## Abstract

Temperature and water availability significantly influence mosquito population dynamics. We’ve devised a method, integrating experimental data with insights from mosquito and thermal biology, to calculate the basic reproduction number (*R_M_*) for urban mosquito species, *Aedes albopictus* and *Aedes aegypti*. *R_M_* represents the number of female mosquitoes produced by one female during her lifespan, indicating suitability for growth. Environmental conditions, including temperature, rainfall and human density influence *R_M_* by altering key mosquito life cycle traits. Validation using data from Spain and Europe confirms the approach’s reliability. Our analysis suggests that temperature increases may not uniformly benefit *Ae. albopictus* proliferation but could boost *Ae. aegypti* expansion. We suggest using vector *R_M_* maps, leveraging climate and environmental data, to predict areas susceptible to invasive mosquito population growth. These maps aid in resource allocation for intervention strategies, supporting effective vector surveillance and management efforts.

## 2 Introduction

Insects manifest intricate life cycles and exhibit rapid responses to environmental variations, posing challenges in the comprehension and tracking of their populations. Some insects, such as dragonflies, mosquitoes, or moths, have life cycles that involve aquatic, terrestrial, and aerial stages, with dynamics shaped by factors such as weather conditions, water availability, landscape features, and human activities (Schowalter 2016). As a result, some insect population dynamics manifest as bursts with highly heterogeneous patterns in both space and time.

Among extensively studied insect species, mosquitoes stand out due to their capacity to transmit global diseases such as malaria and dengue. Modelling insect dynamics, particularly mosquitoes, necessitates a substantial volume of data. Commonly employed data types include presence-absence records and insect trap counts, both incurring significant costs and often inadequately covering the entire inhabited areas of the species. Given these limitations, mathematical approaches become paramount for extrapolating the spatial distribution and temporal evolution of a species.

Numerous mathematical frameworks have been employed to model insect dynamics (Lippi et al. 2023). One notable approach, mostly used for mosquitoes, relies on the principles of thermal biology (Angilletta 2009). This approach explores the relationships between temperature and key traits governing the mosquito’s life cycle(Parham and Michael 2010; Mordecai et al. 2017; Di Pol et al. 2022; Shocket et al. 2020). The thermal biology approach can be integrated with the concept of the basic reproduction number, *R*_0_, (Diekmann et al. 2010) to reveal how mosquito numbers vary in response to temperature fluctuations. In the context of diseases, the basic reproduction number, *R*_0_, is defined as the number of secondary infections in an otherwise susceptible population caused by a single original infection. For the case of mosquitoes (McCormack et al. 2019), the basic reproductive number is also known as the mosquito basic reproduction number, *R_M_*, which defines the number of females mosquitoes produced by a female mosquito in its entire lifespan. Importantly, the application of these frameworks to vector population dynamics remained largely unexplored.

Empirical evidence underscores the pivotal role of temperature in the mosquito life cycle (Delatte et al. 2009; De Majo et al. 2019), temporal dynamics (Marini et al. 2020; Erguler et al. 2016), and spatial distribution (Kraemer et al. 2019). Beyond temperature, mosquito population studies reveal the influence of factors like rainfall (Valdez et al. 2018; Alto and Juliano 2001), human population density (Rodrigues et al. 2015), land cover (Benitez et al. 2020), and relative humidity (Brown et al. 2023; Blanco-Sierra et al. 2023). In this study, we investigate how temperature, rainfall, and human population density modulate *R_M_*, serving as an indicator of mosquito suitability for growth. If *R_M_ >* 1 the female mosquito population experiences exponential growth over time, resulting in an overall population increase. The female mosquito population steadily diminishes until it reaches extinction if *R_M_ <* 1. In other words, *R_M_* determines whether mosquito proliferation may occur after colonizing a specific area. We concentrate on two species of *Aedes* mosquitoes: *Aedes albopictus* and *Aedes aegypti*, potential vectors for diseases like dengue, Zika, Chikungunya or Yellow fever. Notably, *Ae. albopictus* has established populations in Europe since 1979 in Albania (Adhami and Murati 1987) and in Spain since 2004 (Aranda et al. 2006b). *Ae. aegypti* is sporadically arriving to Canary Islands, mainly from Madeira (Portugal) and Cape Verde Islands (Africa), with documented records of adult and larval presence that trigger local surveillance and control campaigns in Spain (ECDC 2018; Centro de Coordinacíon de Alertas y Emergencias Sanitarias 2023), and has also been observed in the Netherlands (Scholte et al. 2010) and Cyprus (Shkurko 2022). To test the validity of the mosquito reproduction number, we analyze the relationship between presence-absence and trap count data in Spain and presence-absence at European level for *Ae. albopictus*. Furthermore, we show the different distributions between the two invasive species *Ae. albopictus* and *Ae. aegypti* and assess the impact of future climate change scenarios on the mosquito reproduction number.

As emphasized by the World Health Organization, the absence of established vaccines underscores the critical role of vector management as a primary strategy in tackling the growing global burden of vectorborne diseases (Balakrishnan 2022). The latter being particularly sensitive to changes in climate and land use (Rocklov et al. 2023). In this context, introducing an innovative suitability index to cast hotspots of vector suitability for growth, therein establishment, can facilitate the implementation of proactive and cost-efficient vector management measures.

## 3 Materials and Methods

### 3.1 Data

To compute the Basic Mosquito Reproduction Number, *R_M_*, we modeled the dynamics of mosquito life cycle parameters modulated by three environmental variables: temperature, rainfall, and human population density. Historical climatic data for 2004 and 2020, including temperature and rainfall, were extracted from the CERRA ERA5 dataset (ECMWF Re-Analysis 5 (Hersbach et al. 2020)). CERRA ERA5 is a freely accessible dataset offering climatic data from 1940 until July 2021, with a temporal resolution of 3h and a spatial resolution of 5.5km. This dataset involves reanalyzed data, combining model outputs with observations using the laws of physics. Human population density data for each year at the municipality scale for Spanish maps was obtained from the INE (Instituto Nacional de Estadıstica), and for Europe, the data was extracted from the fourth version Gridded Population of the World (GPW) collection at 2.5 arc-minute (aproximatly 5 km) for the year 2020 (Columbia University 2018). Additionally, monthly climate data for future predictions for the periods 2041-2060 and 2061-2080 were extracted from the CMIP6 dataset with a resolution of 2.5 minutes of a degree of longitude and latitude. (source: https://worldclim.org/data/cmip6/cmip6climate.html). We used the R package *geodata* (Hijmans et al. 2023) to extract the data. We chose the SSP3-7.0 socioeconomic pathway, which defines the climate change scenario ”Middle of the Road” which represents an intermediate mitigation effort against climate change. We extracted the two variables related to temperature available on the WorldClim website: minimum and maximum monthly average temperature. We then computed mean temperature as the average of these two variables. To mitigate biases originating from specific datasets, all available datasets were gathered and averaged across all sources. These averaged values were then used as representative values for each period and location. For future scenarios, we used population density estimates from the GHS datasets (Freire et al. 2016) for 2030 (source: https://ghsl.jrc.ec. europa.eu/ghs_pop2023.php).

The model was validated using presence-absence and trap count data for *Ae. albopictus* in Spain. Presence-absence data spans the entire invasion process of *Ae. albopictus* in Spain from 2004 up to October 2023. Data from 2004 to 2014 originated from authoritative surveillance at the municipal level, mostly conducted through oviposition and adult traps. From 2014 to 2022, the data combined these authoritative methods with successful citizen science methodologies oriented towards surveillance (Palmer et al. 2017; Bartumeus et al. 2018) (see daily updated maps in Europe at: https://map.mosquitoalert.com/spa/distribution/en). Spatially, the validation involved *Ae. albopictus* presence-absence presence data at the municipal level in five administrative regions of Spain (Andalusia, Aragon, Catalonia, Valencian Community, Basque Country). Also, at the European scale we use presence-absence data for October 2023 from the Mosquito Alert citizen science platform. It combines data from the ECDC agency and citizen science data. Presence-absence data at the European scale corresponds to administrative units at the NUTS 3 level, roughly matching the province level. Spanish municipalities, which are at a smaller scale than NUTS 3, provide presence-absence data at a finer resolution. Therefore we have two data sets with presence-absence, one for Europe and one for Spain with finer resolution. The mosquito trap count data originated from a total of 13 BG-Sentinel traps distributed across various municipalities in the NE Spain (Catalonia region) with the following numbers: Blanes (11), Lloret de Mar (1), Palafolls (1), Tordera (1). The traps operated from November 2020 until December 2022 for Blanes, from September 2020 until December 2022 for Palafolls, from May 2020 until December 2022 for Todera, from June 2020 until November 2020 for Lloret de Mar, and mosquito samples were collected every week.

### 3.2 Vector dynamic model

We have constructed a deterministic compartmental model to capture the dynamics of the mosquito life cycle. The model incorporates three distinct stages: Egg (E), Immature (I), and Adult mosquito (A). Larvae and Pupae stages are amalgamated into the Immature compartment for simplicity. Model parameters, including development rates and mortality rates, depend on temperature (Delatte et al. 2009; Tun-Lin et al. 2000b), while the hatching rate and carrying capacity of the Immature stage are influenced by rainfall and human density. These latter parameters are particularly relevant to the aquatic phases of mosquito development. Rainfall directly affects the availability of natural breeding pools, and human density is presumed to positively correlate with the proliferation and maintenance of water containers resulting from human activities, serving as mosquito breeding sites (Kolimenakis et al. 2021; Rose et al. 2020).

The temporal evolution of each variable is governed by the following equations

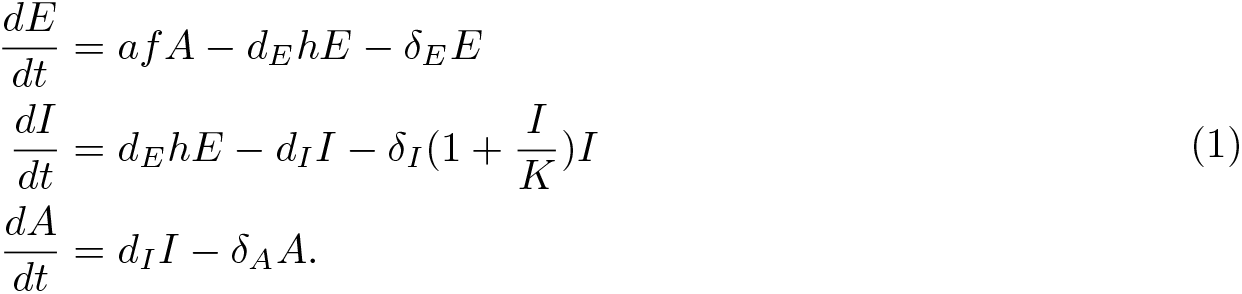

where *δ_X_* are the mortality rate and *d_X_* are the development rates for the *X* compartment. The number of offspring laid by a female mosquito per gonotrophic cycle is denoted by *f*, while *a* defines the biting rate. The proportion of eggs hatching is given by *h* and we define larval resource limitations in the form of a carrying capacity *K*. The underlying assumption is that larvae compete for water availability and resources, being these limiting factors that determine the number of surviving larvae that can transit to the adult stage.

All model parameters depend on temperature, rainfall, and human density (see Table 1). Parameters fall into two groups: one influenced solely by temperature, based on species-specific laboratory data, showing interspecies variations (Mordecai et al. 2017). We establish temperature dependence using methods akin to prior work (Mordecai et al. 2017, 2019), combining thermal biology response functions with laboratory data (Delatte et al. 2009; Calado and Silva 2002a; Dickerson 2007; Calado and Silva 2002b; Juliano et al. 2002; Tun-Lin et al. 2000a; De Majo et al. 2019; Farnesi et al. 2009a; Byttebier et al. 2014; Farnesi et al. 2009b). For some parameters, we obtained the functional form from Mordecai et al. 2017, while we computed the functional form for other parameters that were not available: egg development rates, egg mortality rates and the probability from larva to adult stage for both species (see Supplementary Section 3 for more details). The second group includes parameters tied to human density and rainfall, consistent across species, from literature sources (Metelmann et al. 2019).

**Table 1:**
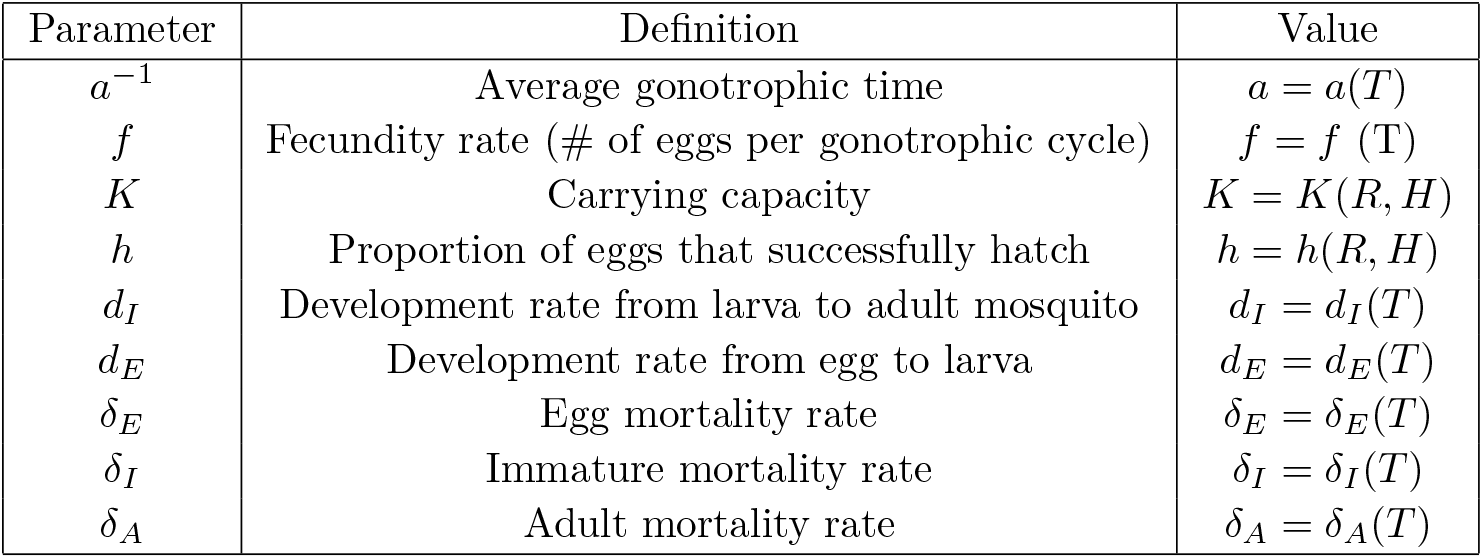
Mosquito model parameters as a function of temperature (*T*), rainfall (*R*), and human density (*H*).

We assume that the functional form for the hatching rate do not differ between *Ae. aegypti* and *Ae. albopictus*. The hatching rate, which describes the proportion of hatching eggs as a function of water container availability, dependent on human density and rainfall, is sourced from (Metelmann et al. 2019) and is given by:

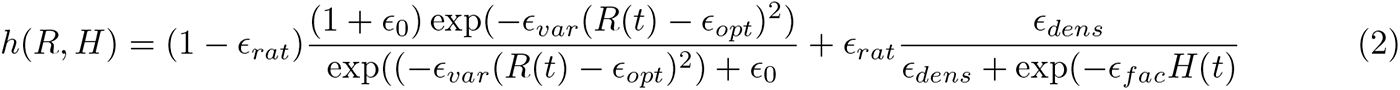

where *H* is the Human density in squared kilometers and *R* is the rainfall in mm. With an optimum at 8 mm of rainfall and a human density above 500 people per square kilometer. Equation 2 defines the proportion of eggs that hatch as a function of rainfall and human density. The first summand represents rainfall impact, following a unimodal pattern; both low and excessive rainfall hinder hatching. The second summand reflects human density influence, saturating at optimal levels. The coefficient *e_rat_* (set at 0.5 for equal weighting) regulates variable importance in hatching rate, specially human density and rainfall.

### 3.3 Mosquito basic reproduction number, ***R_M_***

The mosquito basic reproduction number, *R_M_*, defines the average number of adult female mosquitoes produced by one female mosquito in her entire lifespan (McCormack et al. 2019). Consequently, the value of *R_M_* determines whether the population of female adult mosquitoes will increase or decrease in a given region, conditioned to the existence of an incipient colonizing population.

If *R_M_ >* 1, the population increases and approaches its maximum capacity, limited by the carrying capacity *K*. On the contrary, if *R_M_ <* 1, the population goes extinct, resulting in the absence of adult mosquitoes.

To compute this number, we utilized the method proposed by Diekman et al (Diekmann et al. 2010), defining the Basic Reproduction number as the leading eigenvalue of the Next Generation Matrix, NGM. From the Jacobian matrix of the system (see Supplementary Section 1 for more details), we can compute the Next Generation Matrix *NGM* = *−T* Σ*^−^*^1^,

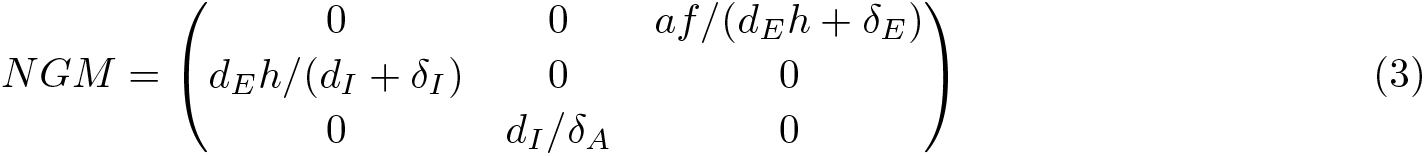

where the leading eigenvalue of this matrix determines the mosquito basic reproduction number,

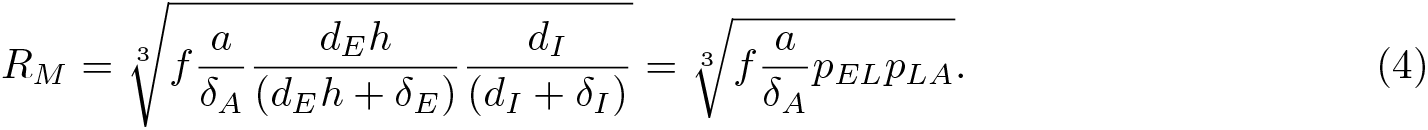

where *p_EL_* defines the probability to develop from egg to larvae and *p_LA_* from larvae to adult. The *R_M_* is the geometric mean of the transition probabilities of each of the populations in the model, with the fecundity rate as an additional factor.

It is important to note that the carrying capacity does not appear in the computation of *R_M_*, which is why we do not explicitly define this parameter. Additionally, the calculation of *R_M_* does not account for transportation factors and assumes that an initial arrival has already occurred in an area with no existing mosquitoes. Therefore, we should consider *R_M_* as a vector suitability index rather than a true invasibility index.

### 3.4 Validation

To validate the suitability index, *R_M_*, we utilized presence-absence data for *Ae. albopictus* in Spain (see Fig. 3c) and Europe (see Supplementary Section 7, Fig. 8). Additionally, we employed count data for *Ae. albopictus* in several cities of Spain, located near the coast at the border between Barcelona and Girona provinces.

The presence-absence map of *Ae. albopictus* in Spain should be understood as a representation of vector establishment as of the year 2023, at the municipality level. This map indicates the initial detections of the invasive mosquito in various municipalities as it spread across the territory over time. It’s important to note that these initial detections should not be interpreted as actual colonization events. Colonization events, which are often linked more to transportation dynamics than climatic factors, are frequently overlooked. Instead, these initial detections are more indicative of the early or potentially delayed establishment of mosquito populations. This establishment process is closely associated with thermo-biological responses, as reflected in the *R_M_* values.

For validation purposes, we aggregated the presence-absence data by region (NUTS2) and computed the percentage of municipalities (the smallest administrative unit) showing established populations of *Ae. albopictus*. Next, we computed the monthly *R_M_* using monthly average temperature and rainfall, along with the annual average human density for the period 2003-2020. This period included one year before the first record of *Ae. albopictus* in Spain (2003) until the last year available in the CERRA ERA5 dataset (2020). Finally, we aggregated municipalities by the number of suitable months (*R_M_ >* 1) for *Ae. albopictus* within the season and assessed, for each suitability value (from 1 to 12 months), the percentage of municipalities with established populations of *Ae.albopictus*.

We also compared weekly mosquito trap counts in different locations with locally computed *R_M_* s for each trap location and week. This was done using the weekly mean temperature and rainfall at ERA5 CERRA cells where the traps were located. For the fitting procedure, we log transformed the trap counts and employed the *lm* function from R (R Core Team 2021). We plotted the results along with the 95% confidence interval.

## 4 Results

### 4.1 Thermo-dependent mosquito basic reproduction number

The mosquito basic reproduction number, *R_M_*, is species-dependent and varies as temperature changes across time and space.. Each of the two *Aedes* species showed a distinct thermal optimum and suitable (i.e., *R_M_ >* 1) temperature range. The differences among species suggested differential reproductive capacities associated with temperature, Fig. 1a. Importantly, in the calculation of *R_M_*, the proportion of hatching eggs depend on water availability. This parameter was determined using Eq. (2) as described in Metelmann et al. 2019, thus being influenced by precipitation and human density, as illustrated in Fig. 1b and Fig. 1c. We applied the same equation for both species, assuming a similar biological response to precipitation and human density. However, it’s worth noting that these environmental variables played a crucial role in delineating the spatial variation in the mosquito basic reproduction number.

**Figure 1:**
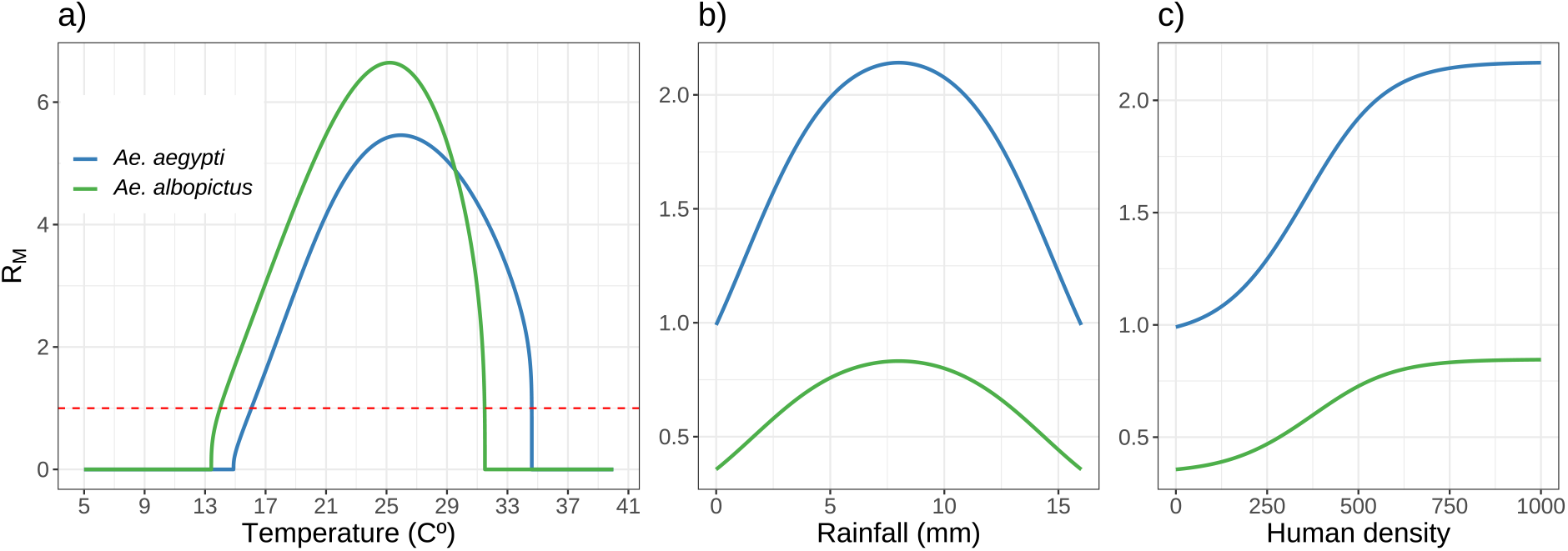
a) The mosquito basic reproduction number, *R_M_*, for the two different species: *Ae. albopictus*, and *Ae. aegypti* as a function of temperature with constant human density (H = 500) and rainfall (R = 8mm). *R_M_* as a function of rainfall (b) and human density (c) with constant temperature (T = 16°C) and H = 0 (b) or R = 0 (c), respectively, to show the effect of each variable. We assumed the same functional form for rainfall and human density for the two species.

By isolating the temperature effect we can better comprehend how temperature uniquely influences each species. We fixed the rainfall and human density to its optima (where the hatching rate, expressed as a proportion, is one, *R* = 8 mm and *H* = 500). For *Ae. albopictus*, the suitable temperature range (where *R_M_ >* 1) is narrower [14.0-31.5 °C] in comparison to *Ae. aegypti*, and the optimum temperature for proliferation is reached at 25.0 degrees (Fig. 1a). *Ae. aegypti* suitability range spans over warmer temperatures [16.1-34.6 °C] and shows an optimum value at 25.9 degrees, higher than the one presented by *Ae. albopictus*. We also show variations in the suitable temperature range (but not in the optimum temperature) for different values of rainfall and human density (Supplementary material Fig. 1 Section 1). Moreover, we conducted a sensitivity analysis to understand the influence of each temperature dependent parameter on the suitability index (see Supplementary Section 4 for more details).

### 4.2 The condition ***R_M_ >* 1** to build Vector Suitability Maps

We can interpret the mosquito basic reproduction number, *R_M_*, as an indicator of the suitability of mosquito population growth, in this case, modulated by temperature, rainfall, and human density. We depicted annual, Fig. 2, and seasonal (Supplementary Material Section 5) vector suitability maps for each targeted species for 2020. To build up both annual and seasonal maps, we used monthly averages of temperature and rainfall and average annual human density. Rather than averaging *R_M_* over months to obtain an annual picture of vector suitability, we followed (Di Pol et al. 2022; Mordecai et al. 2017) and computed the number of months when the vector suitability index is greater than one (*R_M_ >* 1) within a year (Fig. 2). The larger the number of months in a given region, the easier it is for mosquito populations to establish (compared to another region with the same inflow of mosquitoes).

**Figure 2:**
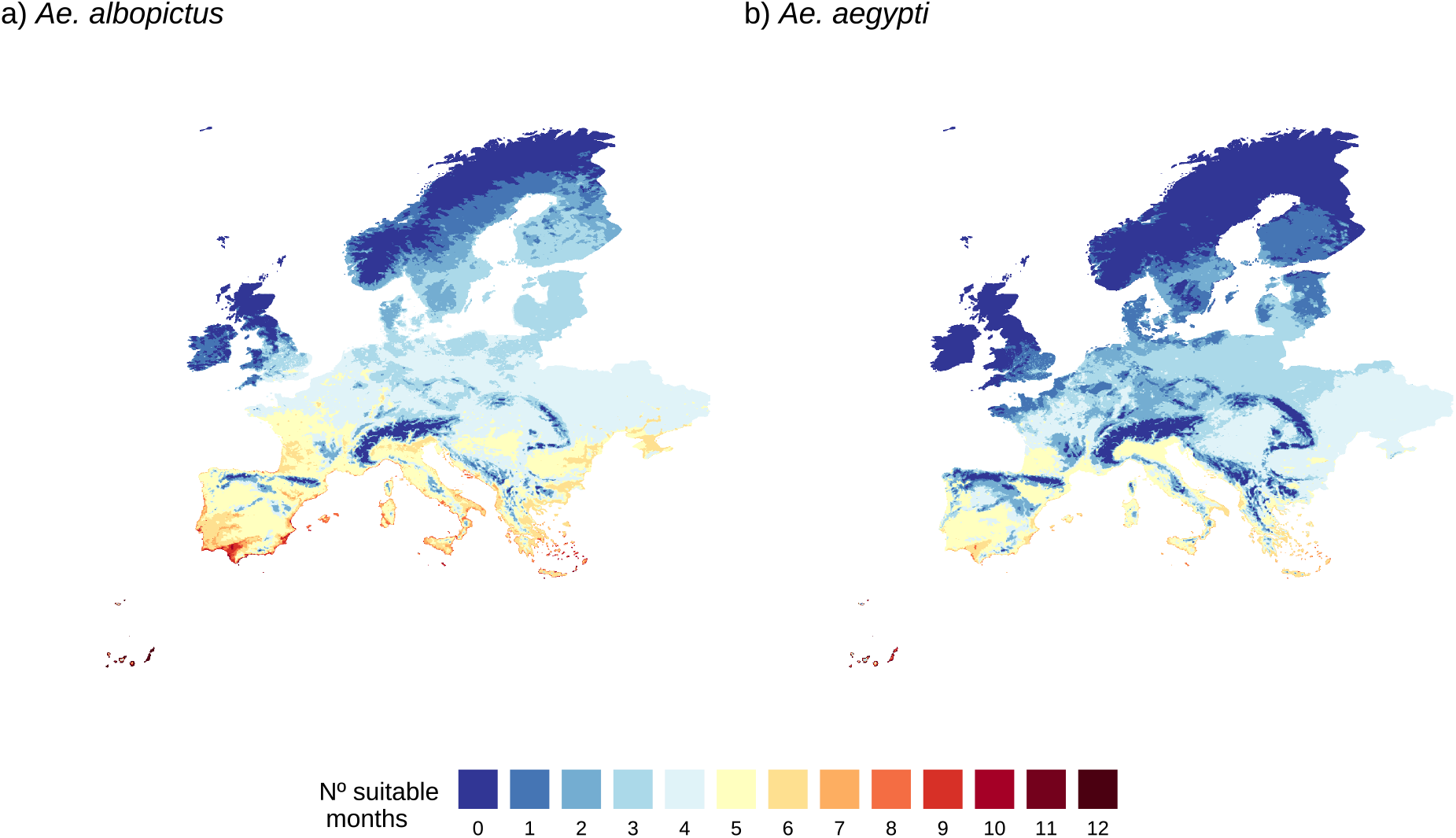
Vector Suitability Maps (Europe 2020) for *Ae. albopictus* (a) and *Ae. aegypti* (b). Suitability is represented as the number of months with *R_M_ >* 1. Color gradient from brown (large number of suitable months) to blue (low number of suitable months).

**Figure 3:**
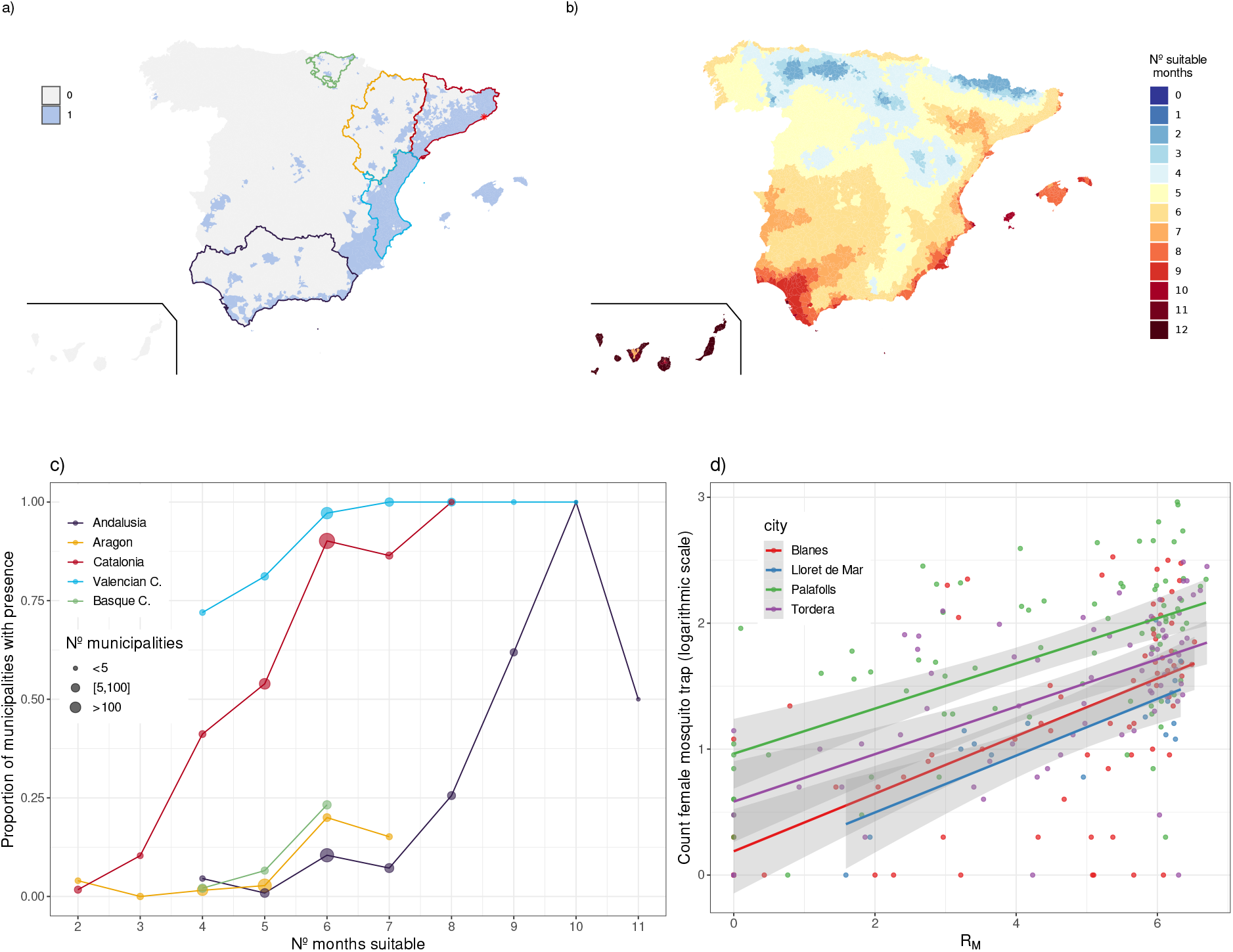
Validation of the mosquito basic reproduction number, *R_M_*, with empirical data. a) Municipality level presence (blue) and absence (grey) of *Ae. albopictus* in Spain in October 2023. b) Number of suitable months (i.e. *R_M_ >* 1) for *Ae. albopictus* in for the period 2003-2020, we compute the *R_M_* with the average rainfall, temperature and human density for this period. c) Relationship between *R_M_* for 6 different regions in Spain and the proportion of municipalities that are positive for the presence of *Ae. albopictus* in the regions. Catalonia and the Valencian Community where *Ae. albopictus* has earlier establishment (first records 2004 and 2005 respectively) compared to the initial presence of these mosquitoes in the Basque Country, Andalusia, and Aragon (2016, 2014 and 2016 respectively). d) Correlations between trap *Ae. albopictus* count data and *R_M_* for four different city locations in the NE Spain (pointed with a star in the map (a): Blanes, Lloret de Mar, Palafolls and Tordera.

*Ae. albopictus* showed higher suitability in Europe compared to *Ae. aegypti*, Fig.2. *Ae. aegypti* demonstrates high suitability in the Mediterranean countries, primarily along the coast, with Spain having the highest suitability for this species. This variation is attributed to the higher thermal minimum presented for *Ae. aegypti* in comparison with *Ae. albopictus*, Fig.1. For the two species, the Mediterranean coast has the highest suitability in almost all of the territory. In addition, the lowest suitability is located in the mountain range (e.g., Pyrenees and Alps) and the northeast European countries. We can see that in the Mediterranean areas, there are some regions where *Ae. albopictus* is suitable for 9 months. This aligns with recent findings where this mosquito was found in traps during winter months in the east and west Mediterranean countries (Lührsen et al. 2023).

### 4.3 Validation with presence-absence and count data of *Ae. albopictus* in Spain

To validate the mosquito basic reproduction number as a vector suitability index, we investigated whether and how the number of suitable months during the period 2003-2020 (i.e. the average *R_M_* for most of the spreading period of *Ae. albopictus* in Spain) is associated with the progressive establishment of local populations at the municipality level (Fig.3a and c). We selected 5 regions (Comunidades Autonomas) at different invasion stages (Fig.3c). In Catalonia and Valencia Community, *Ae. albopictus* is fully established in many municipalities with the first detection of the species in 2004 and 2005, respectively (Aranda et al. 2006a). In Andalusia (south), Aragon (north east, non-coastal) and Basque Country (north) the species was first detected about ten years later: 2014 (Andalusia) and 2016 (Aragon, and Basque Country). As *Ae. aegypti* has not yet colonized Spain, and its current establishment in Europe is extremely limited to a few eastern countries, we did not validate the computation of *R_M_* for this species. In general, the higher the number of suitable months for *Ae. albopictus* the greater the proportion of municipalities with established mosquito populations. In the eastern coastal regions of Spain (Catalonia and Valencian Community), where *Ae. albopictus* first established and spread, over 50% of the municipalities showing an *R_M_ >* 1 for at least 5 months within the season showed established populations of *Ae. albopictus* (Fig. 3c). All municipalities showed full *Ae. albopictus* establishment when *R_M_ >* 1 for at least 8 months within the season. Conversely, in recently colonized areas like Andalusia, Aragon, and the Basque Country, a different pattern emerges as sustained favorable conditions are necessary for full species establishment. Therefore, we only start to see a large proportion of municipalities with established *Ae. albopictus* when *R_M_ >* 1 holds beyond 7 months during the season. Conversely, suitability indexes below 6 months were not associated with *Ae. albopictus* establishments. As an example, municipalities in Andalusia with optimal temperatures (accounting for *R_M_ >* 1) over at least 9 months within the season in order for 50% of them to show *Ae. albopictus* established populations.

A similar analysis at the European scale for *Ae. albopictus* (see Supplementary material Section 7 Fig.8) shows a correlation between suitable months (i.e., months with *R_M_ >* 1) and regions with detected and established populations *Ae. albopictus*.

At the local (city) scale, we validated the suitability index, *R_M_*, with BG-trap female mosquito count data (Fig. 3d). Over three seasons (2020-2022) and in four locations (Blanes, Lloret del Mar, Palafolls, and Tordera), we observed a correlation (Fig. 3d) between *R_M_* for *Ae. albopictus* and the number of mosquitoes captured in traps weekly. *R_M_* was computed using the average temperature and rainfall for each mosquito collection week. As *R_M_* increases for one week, so does the number of *Ae. albopictus* mosquitoes in the trap.

### 4.4 Climate change scenarios for *Aedes* invasive species in Spain and Europe

We examined the suitability index, *R_M_*, for future climate projections across Europe for both *Ae. albopictus* and *Ae. aegypti*. It covers the periods 2041-2060 (Supplementary Fig. 7) and 2061-2080 (Fig.5) For population density, we have used estimates for 2030 (Freire et al. 2016). We also specifically analyzed the suitability index for *Ae. albopictus* at the Spanish level (Fig. 4). Our comparisons included four different time periods: 2004, marking the first recorded presence of *Ae. albopictus* in NE Spain (Catalonia); 2020, the last year covered by the CERRA dataset; and two future climate projections for the periods 2041-2060 and 2061-2080 (Fig.4c,d).

**Figure 4:**
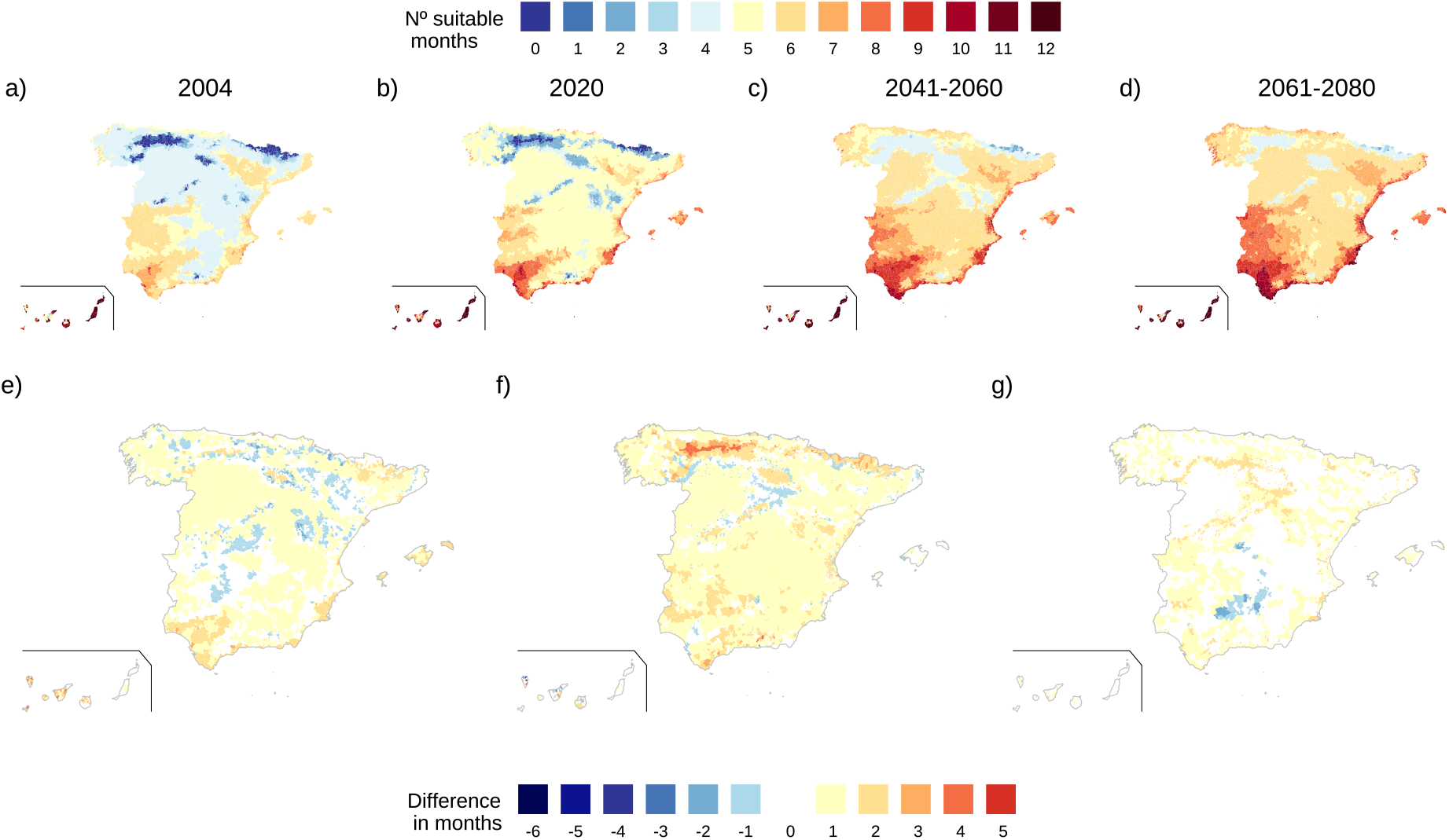
Vector Suitability maps of *Ae. albopictus* in Spain for: 2004 (a), 2020 (b), and two climate future projections from CMIP6 dataset for the periods: 2041-2060 (c), and 2061-2080 (d). Maps comparing different suitability maps and showing in colors the difference in suitable months: from 2020 to 2004 (e), from 2041-2060 to 2020 (f) and 2061-2080 to 2041-2060 (g). Blue colors indicate a decrease in suitability, while brown denotes an increase.

**Figure 5:**
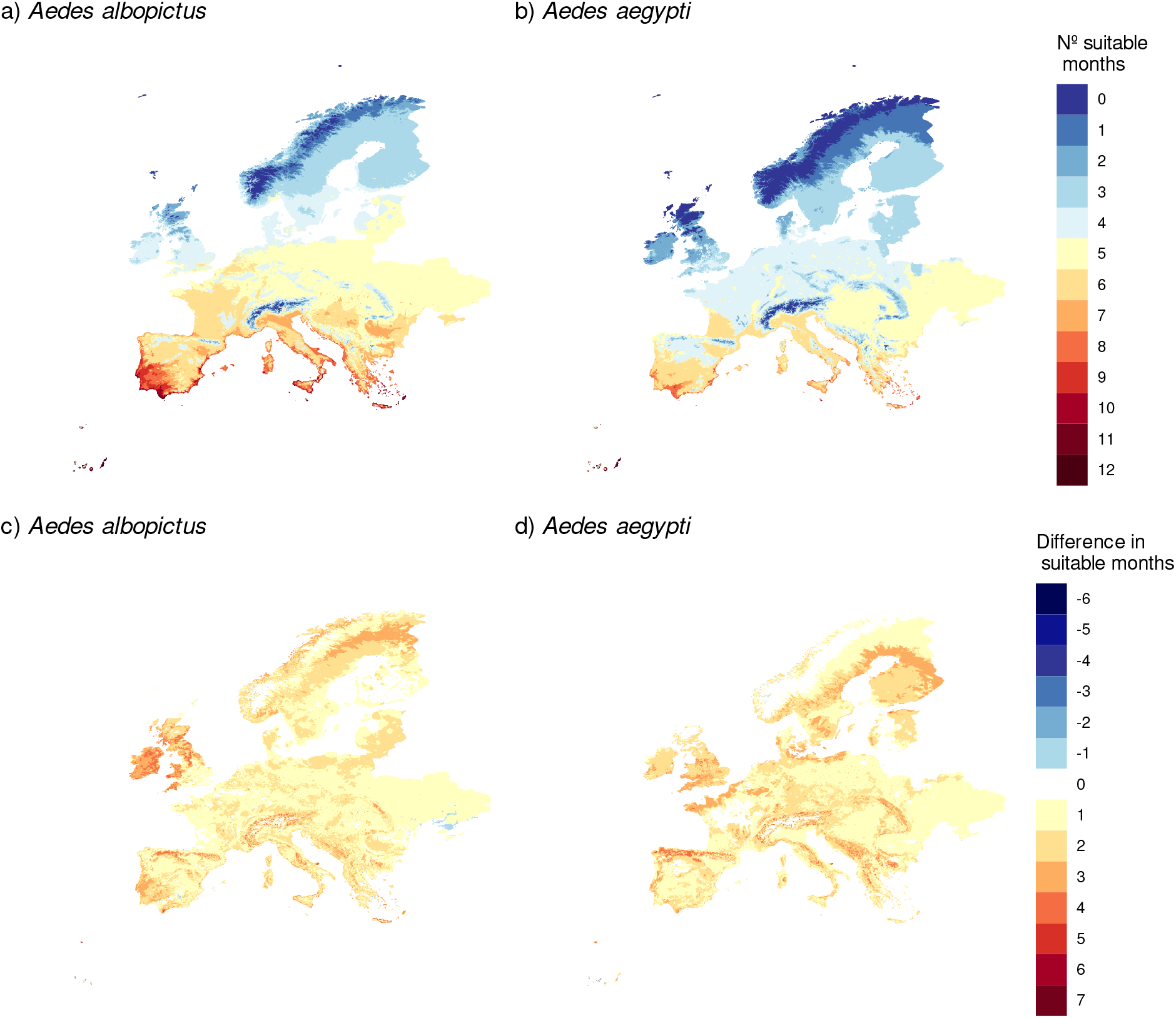
Vector Suitability maps for future climate projections in Europe for the period of 2061-2080 for: *Ae. albopictus* (a) and *Ae. aegypti* (b). Maps showing the difference in suitable months from the period of 2061-2080 and 2020 for : *Ae. albopictus* (c) and *Ae. aegypti* (d).

The number of suitable months where *R_M_ >* 1 in Spain rised from 2004 to 2020 and beyond. Notably, *Ae. albopictus* invaded the NE Spain (Catalonia) in 2004, coinciding with high suitability in the region (Fig. 4a). In 2020 the number of suitable months also increased in the south of Spain (Andalusia) and in the in some coastal regions of the northwest (Galicia) (Fig. 4e), coinciding with the more recent invasion process in the south and a first established population in Galicia in 2023. Overall, Spain shows increased suitability for both periods 2041-2060 and 2061-2080, with significant rises in the northern regions, in particular for the 2041-2060 projection (Fig.4f). Yet, some regions experienced a decline in the number of suitable months from 2004 to 2020 (Fig.4e) due to scarce rainfall and excessive temperatures. In addition, the central-south regions of Spain experienced a decline in the number of suitable months in the projection 2061-2080, attributed also to excessively high summer temperatures exceeding *Ae. albopictus*’ thermal tolerance (Fig.1a).

In Europe, *Ae. albopictus* suitability increased, particularly in southern regions, extending the mosquito activity season. Central and northern Europe showed a comparatively smaller rise in suitable months, while Ireland and England show the most significant increase, with up to four months of suitability. For 2061-2080, the northeast part of Europe showed a projected increase of up to 5 favorable months for *Ae. albopictus*, notably in Scandinavia’s southern regions, deviating from the general trend. This climate change scenario may lead to the establishment of *Ae. albopictus* if migration from already colonized areas occurs. Regarding *Ae. aegypti*, we observed an increase in suitability in southern regions of Europe (Fig.5d), while the northern parts retained fewer suitable months. For certain north-central European regions suitability is bounded to only 4 months during the period 2061-2080 (Fig.5b). Northern France showed a more significant increase, with up to a 4-month rise in suitability, while central France maintained the same level of suitability. Moreover, there was a decrease in suitability resulting from reduced rainfall, despite an increase in temperature, observed in specific regions of the Scandinavian countries and the Mediterranean islands (Fig.5d).

Furthermore, as previously noted, we observed a clear decrease on the number of suitable months for *Ae. albopictus* in Southern Spain, and also in southern Greece, attributed to higher summer temperatures (Fig.5a and Supplementary Fig. 6). The non-linear behavior of the suitability index, *R_M_*, was more apparent in *Ae. albopictus* than in *Ae. aegypti* due to the lower thermal maximum of the former (Fig. 1a). Since *Ae. aegypti* exhibited a *R_M_ >* 1 at higher temperatures when compared to *Ae. albopictus*, potentially leading to increased suitability for this species as summer temperatures rise due to climate change in some areas where *Ae. albopictus* population growth would decrease the number of suitable months.

## 5 Discussion

Mechanistic models grounded on thermo-biology insights have frequently been utilized to calculate the basic reproduction number (*R*_0_) of vector-borne diseases (Mordecai et al. 2019; Di Pol et al. 2022). Yet, their use in evaluating the dynamics of vector populations themselves has been relatively limited (Metelmann et al. 2019), with statistical models typically being employed for this purpose.

Statistical suitability models infer the likelihood of observing a vector, relying on resource-intensive occurrence data (Kraemer et al. 2019; Cunze et al. 2016b,a). Conversely, a suitability framework based on the basic reproductive number (Diekmann et al. (2010)) offers a more precise delineation of temperature ranges and optima, directly depicting the potential for vector population growth under specific environmental conditions. The advantages of computing *R_M_* over other vector suitability models lie in its conceptual simplicity, rapid scalability, and potential for refinement over time, particularly through enhancing thermal-response functions in laboratory settings.

The limitation of the *R_M_* computation becomes evident in the reliance on fixed-temperature laboratory experiments to estimate life cycle trait responses, which lack representation of natural temperature fluctuations. Recent studies aim to address this limitation by exploring temperature fluctuation impacts (De Majo et al. 2019). Additionally, considering other factors influencing vector life cycles, such as rainfall, relative humidity (Brown et al. 2023), human density, and landscape features (Benitez et al. 2020), may contribute to more realistic depictions of basic reproductive numbers and population dynamics. Building on Metelmann et al. (2019), we used rainfall and human density as proxies for mosquito breeding site proliferation, impacting key life cycle parameters of *Aedes* (Li et al. 2015; Benitez et al. 2020), like the egg hatching rate and the carrying capacity. Importantly, since the *R_M_* describes the onset of the next generation from a small initial one, only the egg hatching rate plays a role whereas the carrying capacity vanishes from the *R_M_* computation. Of note, in our model, human density does not influence biting rates but only breeding site availability, which can fluctuate seasonally and geographically.

Our suitability maps show extensive favorable months for *Ae. albopictus* along the Mediterranean coast, aligning with recent findings of winter activity of this species (Lührsen et al. 2023) in Mediterranean regions. More generally, the current temperatures in Europe appear to be more favorable for *Ae. albopictus* compared to *Ae. aegypti*, with the latter exhibiting a higher temperature optimum than the former. This thermal suboptimal scenario may hinder the establishment of *Ae. aegypti* in Europe (Win 2022). Comparing current decades (2020) with 2061-2080, most of Europe may see at least one-month rise in suitability for both species, except parts of Scandinavia. Conversely, mountainous areas like the Alps or Pyrenees could experience a four to seven-month increase. Northwestern countries like England and Ireland may also see a four-month rise of suitability for *Ae. albopictus*, and up to three months for *Ae. aegypti*. However, some coastal Mediterranean regions may face up to a six-month decrease in suitability due to temperatures surpassing the thermal limit for *Ae. albopictus* (Caminade et al. 2012).

Overall, climate change may facilitate the establishment of *Ae. albopictus* in northern Europe and *Ae. aegypti* in southern Europe. While temperature rise generally favors these species, a substantial increase in summer temperatures could impact their suitability (Couper et al. 2021). The temperature increase may result in unsuitable conditions during summer months, potentially disrupting the mosquito season and affecting mosquito dynamics by reducing their abundance or even leading to extinction, as they may struggle to thrive after such a drastic temperature surge. *Ae. aegypti* might be less sensitive to the impact of summer heat waves compared to *Ae. albopictus* due to its higher thermo-tolerance. Additionally, decreased rainfall in the future could further threaten both mosquitoes species by diminishing breeding sites.

When evaluating potential distributions of invasive species, it is crucial to acknowledge their high behavioral plasticity (Juliano and Lounibos 2016). For example, *Aedes* urban species have rapidly adapted to breed in artificial containers produced by humans. More specifically, *Ae. albopictus* has exhibited remarkable adaptability to new environmental conditions (Medley et al. 2019; Powell and Tabachnick 2013), including car-mediated dispersal by adults (Eritja et al. 2017). In the case of mosquitoes, rapid evolution (Papa et al. 2017; Love et al. 2023) is a prominent factor, so that these type of behaviours are rapidly fixed and spread over the populations. All in all, it is evident that future suitability and spatial distribution cannot be exclusively predicted by environmentally-driven population dynamics, but must also consider the adaptive responses of mosquitoes within ever-expanding humanized landscapes.

While recognizing the limitations of basic vector indicators assessing growth suitability, we suggest *R_M_* maps can prove a practical strategy for vector surveillance and proactive measures in regions vulnerable to urban *Aedes* mosquito proliferation.

## Data and Code Availability

The data supporting the results are open and accessible through dedicated web pages and from other literature cited in the main manuscript, except for two Spanish datasets: (i) *Ae. albopictus* presenceabsence data at the municipality level will soon be submitted to a journal. In the meantime, the data can be accessed upon request, and (ii) *Ae. albopictus* trap count data from the period 2018-2022 in Spain uploaded to the same repository as the code. R code used for processing the data and computing the suitability maps is available in the GitHub repository: https://github.com/mpardo1/RM_mosquito.

## Authorship statement

MPA, DA and FB conceived the idea for the study and designed the analyses. RE processed the mosquito presence-absence data. MPA processed the climate data and performed the analyses. MPA wrote the first draft of the manuscript. All authors contributed to manuscript revisions and gave approval for publication.

## Acknowledgements

We acknowledge the World Climate Research Programme, which, through its Working Group on Coupled Modelling, coordinated and promoted CMIP6. We thank the climate modeling groups for producing and making available their model output, the Earth System Grid Federation (ESGF) for archiving the data and providing access, and the multiple funding agencies who support CMIP6 and ESGF. Thanks to Rachel A. Taylor for her advice and feedback about the thermal responses fitting. Also, to Rachel Lowe, Jaume Ramon-Gamon and Alba Llabres to help us to find the more suited datasets for the climatic data. The project leading to these results has received funding from “la Caixa” Foundation (ID 100010434), under agreement HR-18-0036, and the program EU Horizon 2020, Grant agreement “VEO” No. 874735.

