## Supplementary information for "Present and future suitability of invasive and urban vectors through an environmentally-driven Mosquito Reproduction Number"

#### 1 Next Generation Matrix

The mosquito model is given by

$$\begin{aligned}\frac{dE}{dt} &= afA - d_E hE - \delta_E E \\ \frac{dI}{dt} &= d_E hE - d_I I - \delta_I (1 + \frac{I}{K}) I \\ \frac{dA}{dt} &= d_I I - \delta_A A.\end{aligned} \tag{1}$$

where all parameters are described in Table 1. The mosquito basic reproduction number, $\mathcal{R}_M$, for the mosquito model can be computed by means of the NGM, (Diekmann et al. 2010). Decomposing the Jacobian, J, evaluated at the extinction equilibrium of system Eq.(1) as

$$J = \begin{pmatrix} -(hd_E + \delta_E) & 0 & af \\ hd_E & -(d_I + \delta_I) & 0 \\ 0 & d_I & -\delta_A \end{pmatrix} = T + \Sigma = \tag{2}$$

$$= \begin{pmatrix} 0 & 0 & af \\ hd_E & 0 & 0 \\ 0 & d_I & 0 \end{pmatrix} + \begin{pmatrix} -(hd_E + \delta_E) & 0 & 0 \\ 0 & -(d_I + \delta_I) & 0 \\ 0 & 0 & -\delta_A \end{pmatrix}, \tag{3}$$

we can compute the Next Generation Matrix $K = -T\Sigma^{-1}$,

$$K = \begin{pmatrix} 0 & 0 & af/(hd_E + \delta_E) \\ hd_E/(d_I + \delta_I) & 0 & 0 \\ 0 & d_I/\delta_A & 0 \end{pmatrix} \tag{4}$$

obtaining the following expression for the $\mathcal{R}_0$

$$\mathcal{R}_0 = \sqrt[3]{f \frac{a}{\delta_A} \frac{d_I}{(d_I + \delta_I)} \frac{hd_E}{(hd_E + \delta_E)}} = \sqrt[3]{f \frac{a}{\delta_A} p_{EL} p_{LA}}. \tag{5}$$

The influence of each variable in the mosquito basic reproduction number, $R_M$, for each specie is shown in detail in Fig.1. Both species narrow their suitable range as rainfall decreases from the optimal

| Parameter | Definition | Value |
|---|---|---|
| $a^{-1}$ | Average gonotrophic time | $a = a(T)$ |
| $f$ | Fecundity rate (# of eggs per gonotrophic cycle) | $f = f$ (T) |
| $K$ | Carrying capacity | $K = K(R, H)$ |
| $h$ | Proportion of eggs that successfully hatch | $h = h(R, H)$ |
| $d_I$ | Development rate from larva to adult mosquito | $d_I = d_I(T)$ |
| $d_E$ | Development rate from egg to larva | $d_E = d_E(T)$ |
| $\delta_E$ | Egg mortality rate | $\delta_E = \delta_E(T)$ |
| $\delta_I$ | Immature mortality rate | $\delta_I = \delta_I(T)$ |
| $\delta_A$ | Adult mortality rate | $\delta_A = \delta_A(T)$ |

Table 1: Mosquito model parameters as a function of temperature ($T$), rainfall ($R$), and human density ($H$).

8mm. The same behavior is observed for lower human density, specifically below 500 people per square meter. *Ae. albopictus* shows a narrower range but similar maximum for $R_M$ in comparison with *Ae. aegypti*.

### 2 Equilibria and feasibility

There are two equilibrium points in the mosquito model: the extinction equilibrium point, where there is no mosquito population, and the non-trivial equilibrium point $P^* = (E^*, I^*, A^*)$, where:

$$E^* = \frac{af}{(hd_E + \delta_E)} A^* \tag{6}$$

$$I^* = \frac{\delta_A}{d_I} A^* \tag{7}$$

$$A^* = \frac{d_I^2}{\delta_A^2 \delta_I} K \left( \frac{d_E haf}{(d_E h + \delta_E)} - \frac{(\delta_I + d_I)\delta_A}{d_I} \right) \tag{8}$$

$$\tag{9}$$

This equilibrium point will be biologically feasible if all its components are greater than zero. The value for the adult mosquitoes at equilibrium is a positive constant time the value for the larva. Therefore if the $I^* > 0$ the equilibrium point is feasible.

$$\left[ \frac{d_E haf}{(d_E h + \delta_E)} - \frac{(\delta_I + d_I)\delta_A}{d_I} \right] > 0 \tag{10}$$

which is equivalent to

$$\frac{af d_E h d_I}{\delta_A (d_I + \delta_I)(d_E h + \delta_E)} > 1 \tag{11}$$

therefore, the same condition holds for the existence of the non-trivial equilibrium point as for the one given by the basic mosquito reproduction number, Eq. (5). If a number is greater (or less) than one, its cubic square root is also greater (or less) than one.

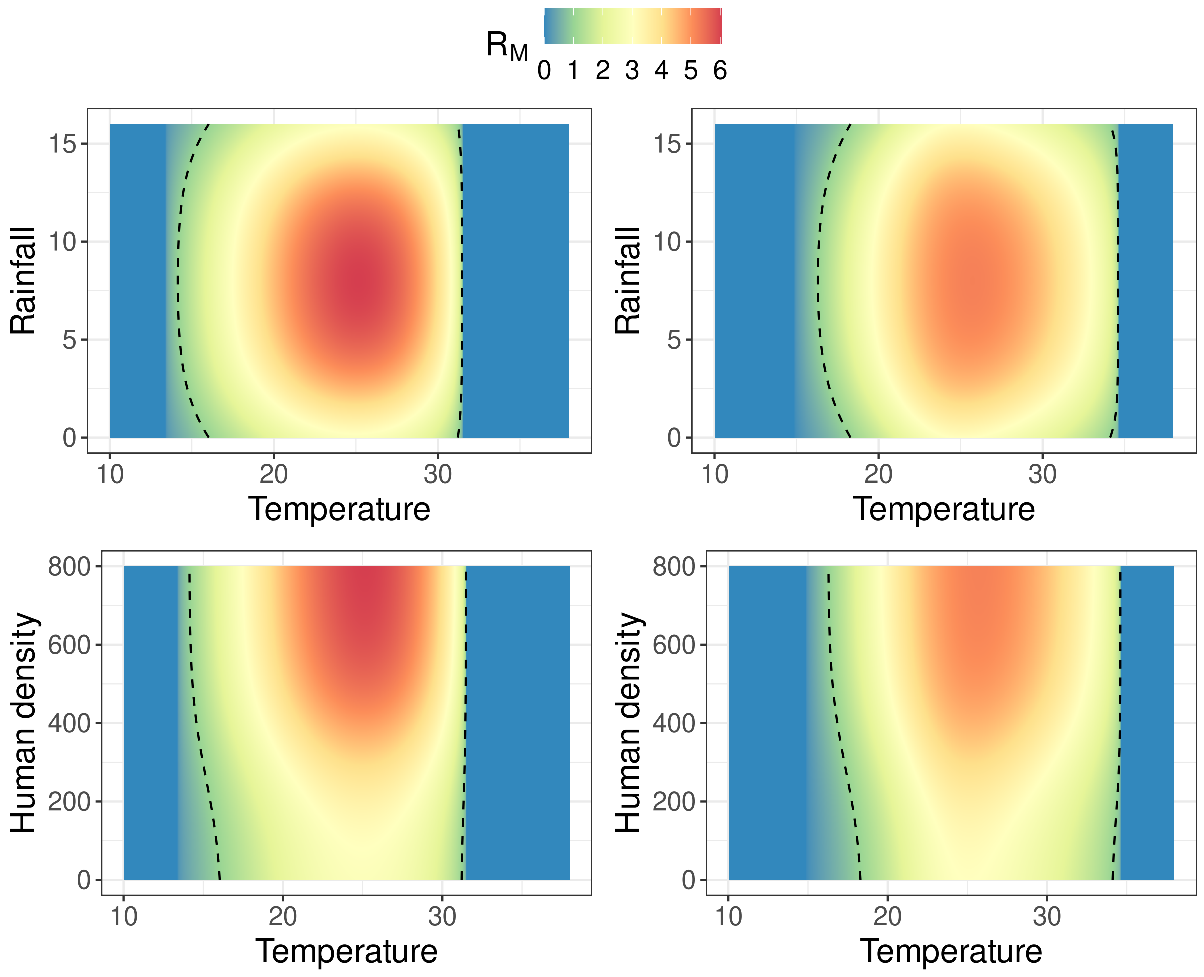


Figure 1: Influence of rainfall (mm) and temperature ($^{\circ}C$) (top) and human density (Number of individuals per squared kilometer) and temperature (bottom) on the mosquito basic reproduction number, $R_M$, for the two species: *Aedes albocpitus* (left) and *Aedes aegypti* (right). Dashed line represents $R_M = 1$. When one variable in a plot its not shown, e.g. top left figure the human density, this variable is set to zero. To isolate the effect of the other two variables. Red colors indicate higher suitability index, $R_M$, while blue colors define low values of $R_M$.

### 3 Thermal responses

To compute the basic reproductive number, we require functional forms linking each life cycle trait to temperature. Some were taken from Mordecai et al. 2017, while others were derived in this study. The exact forms for adult lifespan, $lf$ (inverse of the adult mortality rate, $\delta_A = 1/lf$), and biting rate for *Ae. aegypti* and *Ae. albopictus* were sourced from Mordecai et al. 2017. Additionally, for *Ae. albopictus*, fecundity ($f$), number of eggs per gonotrophic cycle, was obtained from Mordecai et al. (2017). For *Ae. aegypti*, although no direct function for fecundity ($f$) is provided in Mordecai et al. 2017, we utilized the number of eggs laid per day as reported in their study. In Equation 5, the number of eggs laid per day can be calculated by multiplying the inverse of the gonotrophic cycle length (denoted as $a$) by the fecundity ($f$), represented as $fa$.

For the rest of life cycle traits that were not available in Mordecai et al. 2017 we inferred new functions with data from the literature (Delatte et al. 2009; Calado and Silva 2002a; Dickerson 2007; Calado and Silva 2002b; Juliano et al. 2002; Tun-Lin et al. 2000; De Majo et al. 2019; Farnesi et al. 2009a; Byttebier et al. 2014; Farnesi et al. 2009b). We fitted the following parameters: egg development rates, $d_E$, egg mortality rates, $\delta_E$, and the probability from larva to adult, $p_{LA}$, for both *Ae. albopictus* and *Ae. aegypti*, table 2.

The thermal responses are modeled with three different functional forms depending on the data:

- Quadratic $f(T) = -c(T - T_{min})(T - T_{max})$
- Brière $f(T) = cT(T - T_{min})(T_{max} - T)^{1/2}$
- Normal quadratic $f(T) = cT^2 + T_{min}T^2 + T_m axT$

We have used the same notation for the normal quadratic form as for the other two for simplicity. However, these values, $T_{min}$ and $T_{max}$, do not have a straightforward meaning beyond the meaning of the parameters of a parabola.

The values fitted with the nls function in R (R Core Team 2021) are given in table 2. In order to understand if non-linear thermal response models were better than simple linear models, we computed the AIC values for both of them and selected the best model as the one with lower AIC. Table 3 shows different values of AIC for all the parameters fitted in this work. The computation of the probability from larva to adult involved population percentages (by experiment and temperature), resulting in estimates with fewer data points compared to developmental rates, data at individual level, Fig. 2. The probability from larvae to adult mosquito for *Ae. albopictus* presented a lower optimum temperature, 22.8ºC, in comparison with *Ae. aegypti* at 24.8ºC, 2. The egg development rate, $d_E$, for *Ae. aegypti*, showed a higher optimum temperature, 27.3ºC, while *Ae. albopictus* the optimum is reached at 25.5ºC, 2. This parameter exhibited a significant difference between the two species. The egg mortality rates, $\delta_E$, showed a very similar minima for the two species, with an optimum of 23.7ºC and 23.5ºC for *Ae. albopictus* and *Ae. aegypti*, respectively, Fig. 2.

### 4 Sensitivity Analysis

We conducted a sensitivity analysis in order to understand the extent to which parameters change as temperature changes, contributing to variation of $R_M$. This sensitivity analysis involves isolating the con-

| **Parameter** | **Specie** | **Refs** | **Function** | **c** | | $t_{min}$ | | $t_{max}$ | |
|---|---|---|---|---|---|---|---|---|---|
| | | | | mean | sd | mean | sd | mean | sd |
| $p_{IA}$ | *Ae. albopictus* | (Delatte et al. 2009) | Quad | 2.66e-03 | 6.20e-04 | 6.66 | 1.65 | 38.92 | 1.75 |
| $d_E$ | *Ae. albopictus* | (Calado and Silva 2002a) | Quad | 7.12e-04 | 1.11e-04 | 1.7 | 1.26 | 40.51 | 1.51 |
| $\delta_E$ | *Ae. albopictus* | (Dickerson 2007)<br>(Calado and Silva 2002b)<br>(Juliano et al. 2002) | QuadN | 1.93e-03 | 6.18e-04 | -9.1e-02 | 0.03e-02 | 1.33 | 0.36 |
| $p_{IA}$ | *Ae. aegypti* | (Tun-Lin et al. 2000)<br>(De Majo et al. 2019) | Quad | 4.18e-03 | 8.36e-04 | 9.37 | 1.19 | 40.26 | 1.48 |
| $d_E$ | *Ae. aegypti* | (Farnesi et al. 2009a) | Bri | 3.77e-04 | 3.15e-05 | 14.88 | 7.39e-01 | 37.42 | 3.87e-01 |
| $\delta_E$ | *Ae. aegypti* | (Juliano et al. 2002)<br>(Byttebier et al. 2014)<br>(Farnesi et al. 2009b) | QuadN | 4.47e-03 | 5.49e-04 | -2.10e-01 | 2.86e-02 | 2.55 | 3.36e-01 |

Table 2: Table with parameter values fitted in this work by non-linear least-squares (nls) to obtain the thermal responses for each species. The parameters that do not appear in this table come from Mordecai et al (2017) (Mordecai et al. 2017).

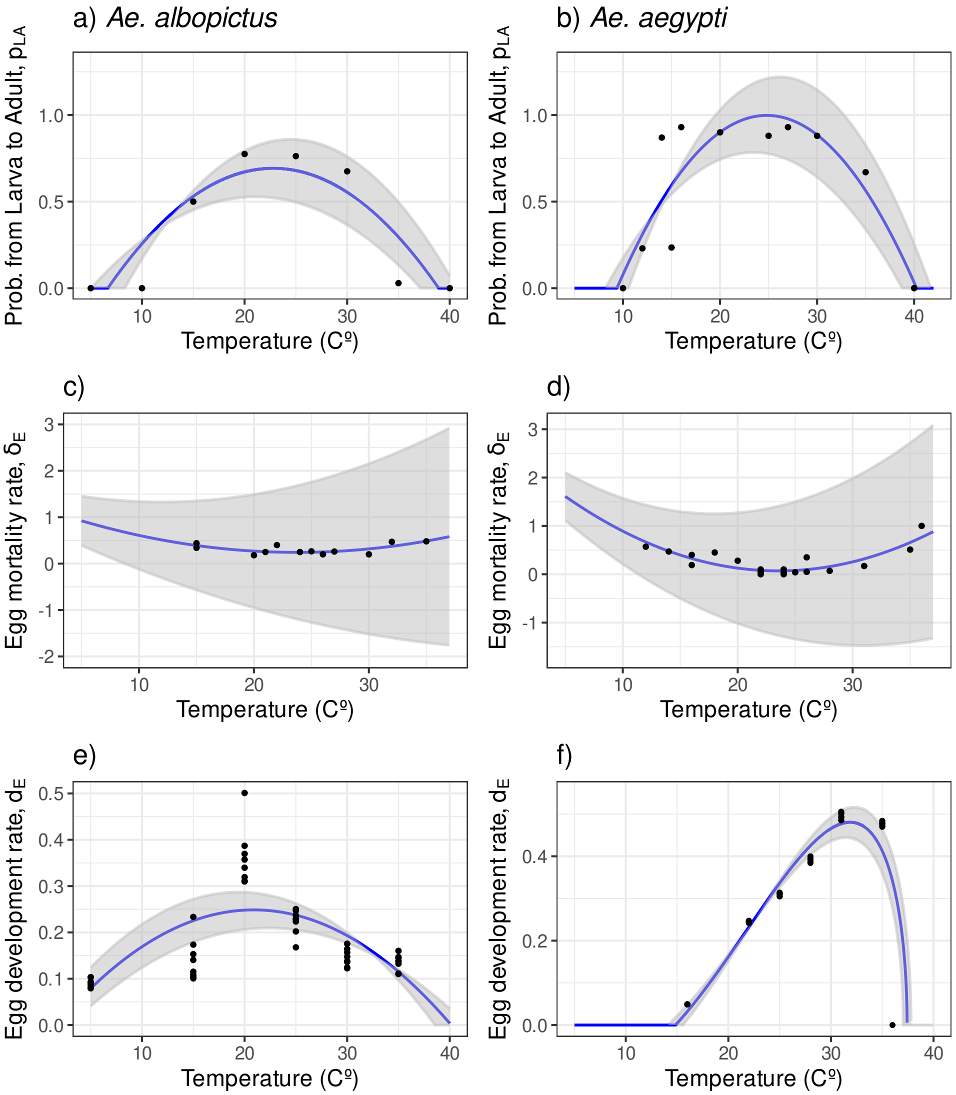


Figure 2: Thermal responses for *Ae. albopictus* and *Ae. aegypti* with the functional form and parameters given in Table 2. The grey area shows the thermal responses for the mean parameter minus (plus) the standard deviation. While the blue line shows the functional form for the mean parameter values. The wider the bandwidth the higher uncertainty there is on the parameter estimation.

| Parameter | Specie | Model | AIC value |
|---|---|---|---|
| $p_{LA}$ | *Ae. aegypti* | Quad | 1.83 |
| $p_{LA}$ | *Ae. aegypti* | Lin | 15.43 |
| $d_E$ | *Ae. aegypti* | Brie | -128.69 |
| $d_E$ | *Ae. aegypti* | Lin | -94.60 |
| $\delta_E$ | *Ae. aegypti* | QuadN | -21.02 |
| $\delta_E$ | *Ae. aegypti* | Lin | 7.14 |
| $p_{LA}$ | *Ae. albopictus* | Quad | 1.26 |
| $p_{LA}$ | *Ae. albopictus* | Lin | 11.62 |
| $d_E$ | *Ae. albopictus* | Quad | -111.77 |
| $d_E$ | *Ae. albopictus* | Lin | -2.46 |
| $\delta_E$ | *Ae. albopictus* | QuadN | -21.04 |
| $\delta_E$ | *Ae. albopictus* | Lin | -14.23 |

Table 3: AIC for the different models tested to fit the laboratory experimental data for each parameter for both species: *Ae. albopictus* and *Ae. aegypti*.

tribution of each model parameter, Fig.3. We have approached this in two distinct manners. Firstly, we isolate the contribution of each parameter to the derivative of $R_M$ with respect to temperature, $dR_M/dT$. More specifically, we are measuring how temperature-driven variations in each of the parameters contribute to variation in the rate of change of $R_M$. Mathematically, this contribution can be expressed as $(dR_M/dx)(dx/dT)$ for each parameter $x$, Fig.3a,b. If the graph of the derivative, Fig.3a,b, shows a steep increase or decrease at a particular point, it suggests that minor temperature changes can lead to significant fluctuations in the vector suitability index due to variations in that parameter. On the contrary, if there is a small increase or decrease in $dx/dT$, it means that $R_M$ is not very sensitive to changes in the parameter value. Additionally, the peaks establish distinct ranges of sensitivity for each parameter. Secondly, we set each parameter to its maximum value and compute $R_M$, then compare it with the original $R_M$ where all parameters depend on temperature, Fig.3c,d.

In the case of *Aedes albopictus*, the adult mortality rate, $\delta_A$, introduces the most variability in $R_M$, Fig. 3a. Additionally, fecundity (daily number of eggs: $fa$) introduces variability in the magnitude of $R_M$, albeit not within the suitable range ($R_M > 1$), Fig. 3c.

For *Aedes aegypti*, fecundity ($fa$) induces higher variability, particularly at the upper thermal limit, Fig. 3d. The remaining parameters introduce similar variability, Fig. 3b.

### 5 Seasonal variability in Spain

We show the seasonal variability of the suitability index, $R_M$, in Spain for 2020. Monthly maps from March to November, for both *Ae. albopictus*, Fig.5, and *Ae. aegypti*, Fig.4, clearly show the increase in suitability during the summer months. The onset (March) follows a south-to-north pattern and the end (November) an inverse north-to-south pattern onset suitability pattern . These patterns look consistent among the two species.

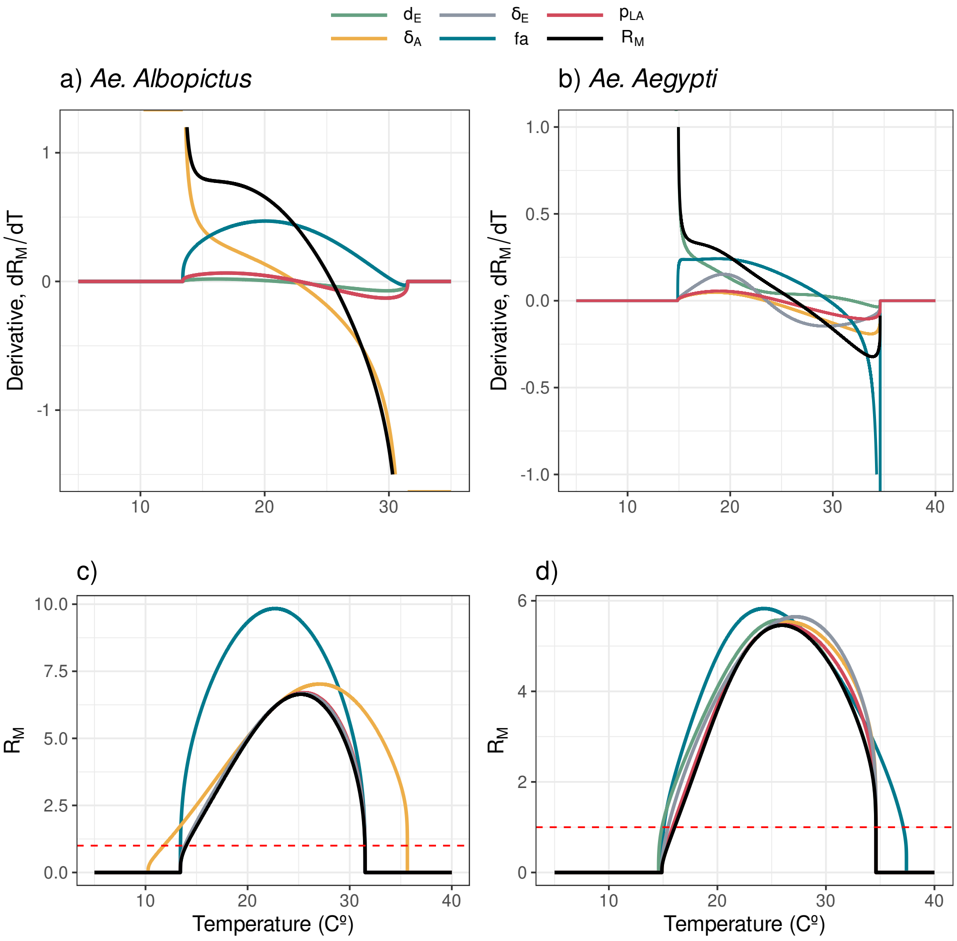


Figure 3: Sensitivity analysis with two methods: calculating the derivative of $R_M$ with respect to each parameter dependent on temperature, denoted as $(dR_M/dx)(dx/dT)$ (top), and $R_M$ with each parameter set to its optimal value (bottom), for each species: Aedes albopictus (left) and Aedes aegypti (right). Parameters analyzed: egg development rate ($d_E$), egg mortality rate ($\delta_E$), probability from larva to adult mosquito ($p_{LA}$), adult mortality rate ($\delta_A$), fecundity, number of eggs per day, ($fa$) and the mosquito basic reproduction number $R_M$.

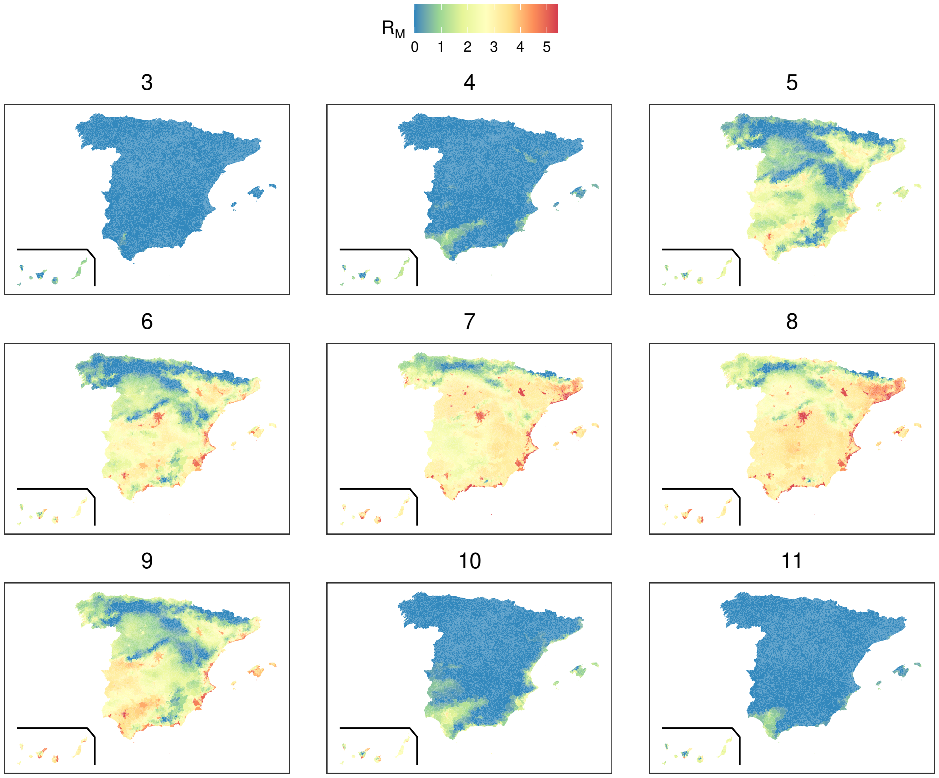


Figure 4: The suitability index, $R_M$ for each month from March to November for *Aedes aegypti* for 2020.

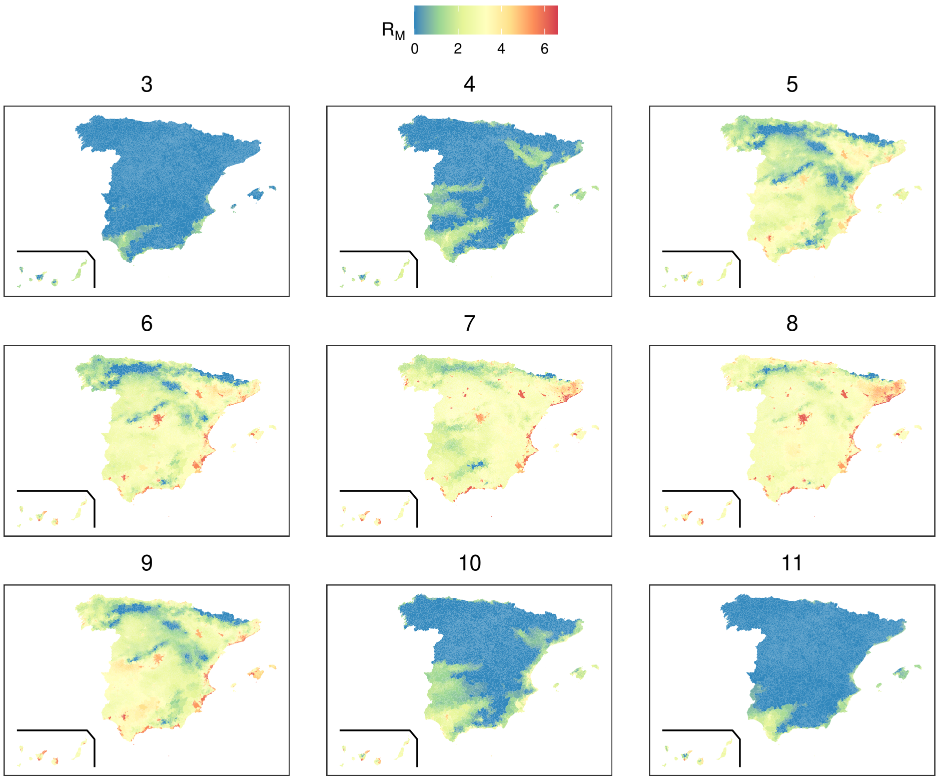


Figure 5: The suitability index, $R_M$ for each month from March to November for *Aedes albopictus* for 2020.

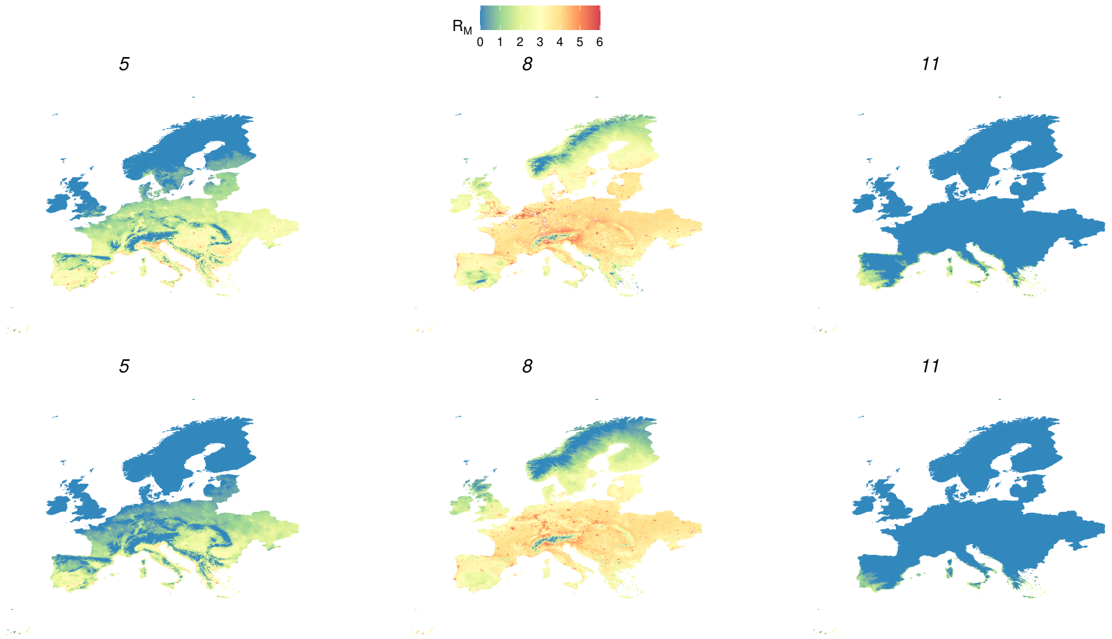


Figure 6: Suitability maps $R_M$ in Europe for the two species: *Ae. albopictus* (top) and *Ae. aegypti* (bottom). Maps are shown for three months of the year: May (5), August (8) and November (11) for the future climate projection 2061-2080.

### 6 European suitability maps future predictions

Here we show the suitability index, $R_M$, of the two species in the period 2041-2060 at a European scale, Fig.7. We note a reduction in suitability for *Ae. albopictus* in Spain's southern interior area during August. This phenomenon is absent in *Ae. aegypti*, attributed to its elevated thermal tolerance. Examining the period 2041-2060 in Fig. 7, we observe an increase in suitability for *Ae. albopictus* across South Europe and an expansive increase in the northern regions, excluding Scandinavian countries, Ireland, and England. Conversely, *Ae. aegypti* does not exhibit a similar surge, with its suitability confined to South Europe. When assessing the differences between period 2041-2060 and period 2061-2080 we can see a decrease in the suitability of *Ae. albopictus* in non-coastal regions of Spain and Greece (blue color) and an increase in western France and other eastern European regions. As for *Ae. aegypti* we observe an increase in eastern regions of Europe both north and centre.
We observe a decrease in suitability for the summer months, in particular in

### 7 Validation *Aedes albopictus*

We have validated the suitability index $R_M$ for *Aedes albopictus* in Europe with presence absence data, Fig.8A. Same procedure as in Section 4.3 main text. We observe a clear relationship between the number of suitable months and the proportion of regions positive for *Aedes albopictus*. We observed a steep increase in the proportion of positive regions reaching forty percent of presence after 5 months of suitability. We also show the supperposition of the suitability map for the period 2003-2020 for *Aedes albopictus* in

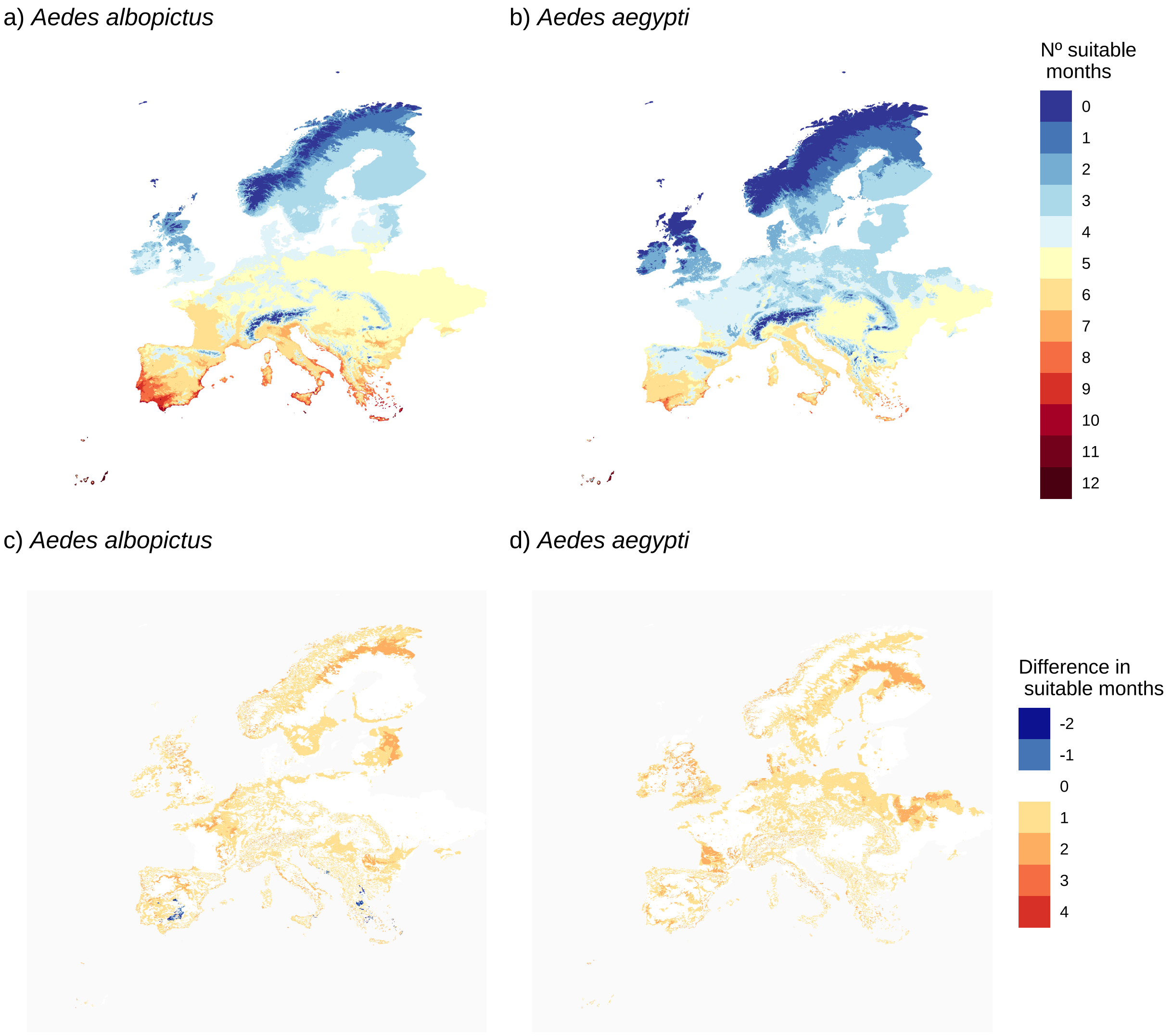


Figure 7: Suitability maps $R_M$ in Europe for the two species: *Ae. albopictus* (a) and *Ae. aegypti* (b) for the period 2041-2060. Differences in the number of suitable months comparing the period 2041-2060 against 2061-2080 for *Ae. albopictus* (c) and *Ae. aegypti* (d).

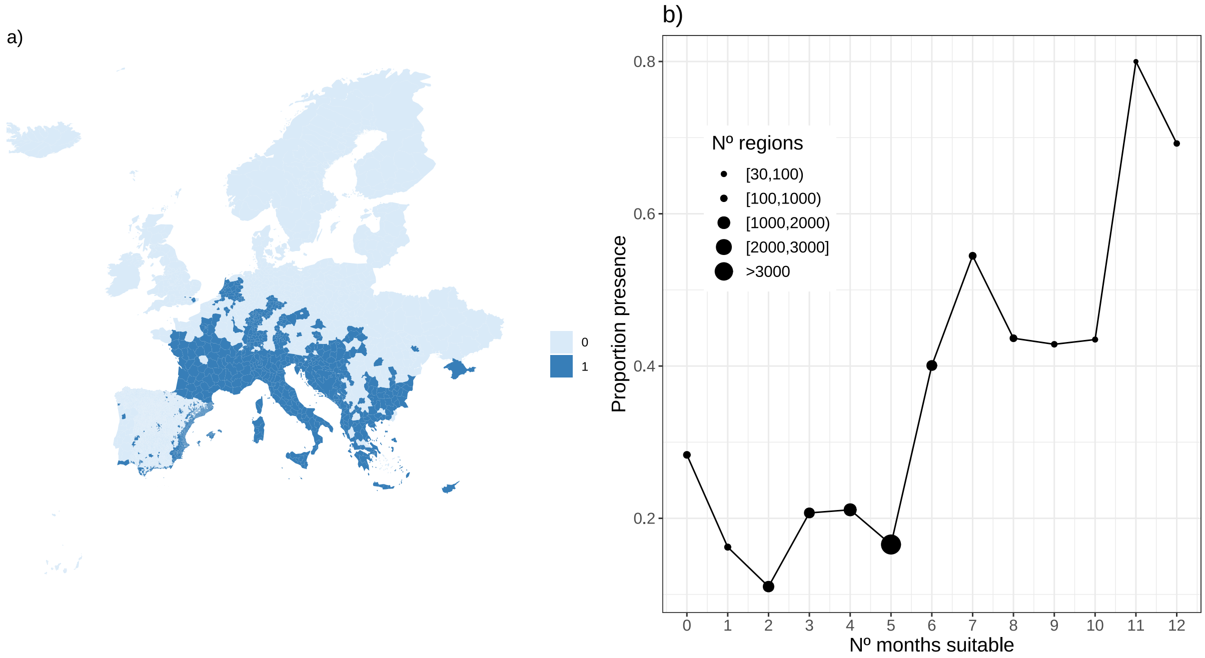


Figure 8: Validation with presence (detection) absence data of *Ae. albopictus* in Europe. a) shows the presence, 1, absence, 0, map for*Ae. albopictus* in Europe. b) shows the proportion of regions NUT3 with presence of *Ae. albopictus* as a function of the number of months suitable.

Spain and the municipalities with presence of *Ae. albocpitus*, Fig.9.

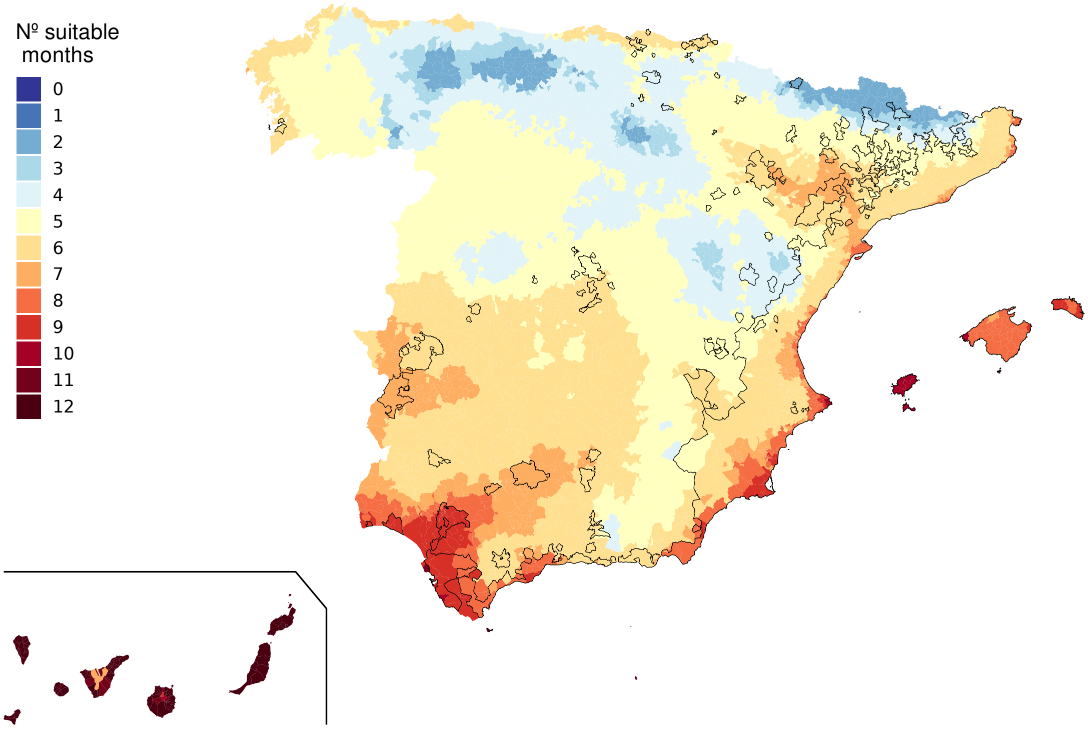


Figure 9: Municipalities with presence of *Aedes albopictus* (delineated regions) and the number of suitable months, $R_M > 1$, for the period 2003-2020 for Spain.
